# Early Invasion Genomic Resources for *Xenia umbellata* (Native to the Red Sea) and its Associated Dinoflagellate Symbionts in Puerto Rico

**DOI:** 10.1101/2025.10.03.680359

**Authors:** Alex J. Veglia, Daniel A. Toledo-Rodriguez, Joyah A. Watkins, Nikolaos V. Schizas

**Affiliations:** Department of Biology, University of Puerto Rico at Mayagüez, PO Box 9000, Mayagüez, PR 00681, USA; EcoAzul, HC01 Box 4729, Lajas, PR 00667, USA; Department of Marine Sciences, University of Puerto Rico at Mayagüez, PO Box 9000, Mayagüez, PR 00681, USA; BioSciences Department, Rice University, 6100 Main St, Houston, TX 77005, USA

**Keywords:** Pulse coral, Caribbean, Hologenome, Symbiodiniaceae, Repeatome, Puerto Rico

## Abstract

The frequency of xeniid soft coral invasions on Caribbean coral reefs is increasing, with three alien species reported so far. *Xenia umbellata* (Anthozoa, Octocorallia, Malacalcyonacea, Xeniidae), native to the Red Sea, was first reported on Puerto Rico coral reefs in October 2023. Here, we present the first draft genome assembly and early-invasion genomic resources for the rapidly spreading *X. umbellata* and its dinoflagellate symbiont (Family Symbiodiniaceae) produced from a specimen collected five months after the initial report. Using deep Illumina metagenomic sequencing (∼243X coverage), we obtained ∼272.4 million high-quality 150 bp reads. The *X. umbellata* draft genome assembly is 151.14 Mbp in length, composed of 27,739 scaffolds, with an N50 of 6,477,837 bp. GenomeScope2 predicted a haploid genome size of 171.6-171.9 Mbp and calculated a heterozygosity of 1.27-1.29%. This suggests that the assembly captures ∼88% of the *X. umbellata* genome, and the relatively high heterozygosity may indicate introduction from a genetically diverse wild population. Completeness was further supported by BUSCO analysis (anthozoa_odb12 lineage), which identified 91.4% complete and 3.9% fragmented BUSCOs. Furthermore, 555,596 sequences were identified as belonging to Symbiodiniaceae, of which 99.97% (n= 555,520) aligned to a *Durusdinium* reference genome, suggesting that the co-invading symbiont belongs to the genus *Durusdinium*. Establishing these genomic and symbiotic resources at an early stage of invasion provides a critical foundation for monitoring range expansion, investigating host–symbiont evolutionary dynamics, and identifying genomic features that may underlie the invasive potential of *X. umbellata* as it spreads across Puerto Rico and the wider Caribbean.

## 1. Introduction

The octocoral *Xenia umbellata* Lamarck, 1816 is a common component of Red Sea coral reef ecosystems, known for its high tolerance to environmental stressors and rapid proliferation, facilitated by whole-body regeneration from even a single tentacle (Halász et al., 2019; Nadir et al., 2023; Mezger et al., 2022). This remarkable regenerative capacity has also established *X. umbellata* as an emerging model system for studying regeneration (Nadir et al., 2023). In October 2023, initial reports from recreational divers documented the onset of a *Xenia umbellata* invasion -- originally misidentified as *Unomia stolonifera*, which is actively spreading in Venezuela (Ruiz-Allais et al., 2014, 2021) -- on Puerto Rico’s already degraded reefs, raising concerns about the species’ potential to take over ecosystems both local and in neighboring islands with little natural resistance (Toledo-Rodriguez et al., 2025). Capable of colonizing available substrate (i.e., coral rubble, rocky substrate, and bare sand) and overgrowing native benthic organisms like ecosystem engineering stony corals and sponges (Figure 1), significant effort from Puerto Rico’s Department of Natural and Environmental Resources Emergency Response Unit has been dedicated to tracking and eradicating *X. umbellata* patches, yet new patches continue to be found in shallow (<30 ft) and deeper reefs (up to 55 m). As the region prepares for the long-term management of *X. umbellata’s* potential impact on Caribbean reefs, a lack of genomic and microbial resources for this species remains. Such genomic data, especially generated during the early phase of the invasion, can provide critical insights into its invasion dynamics and establish a foundation for future research.

**Figure 1.**
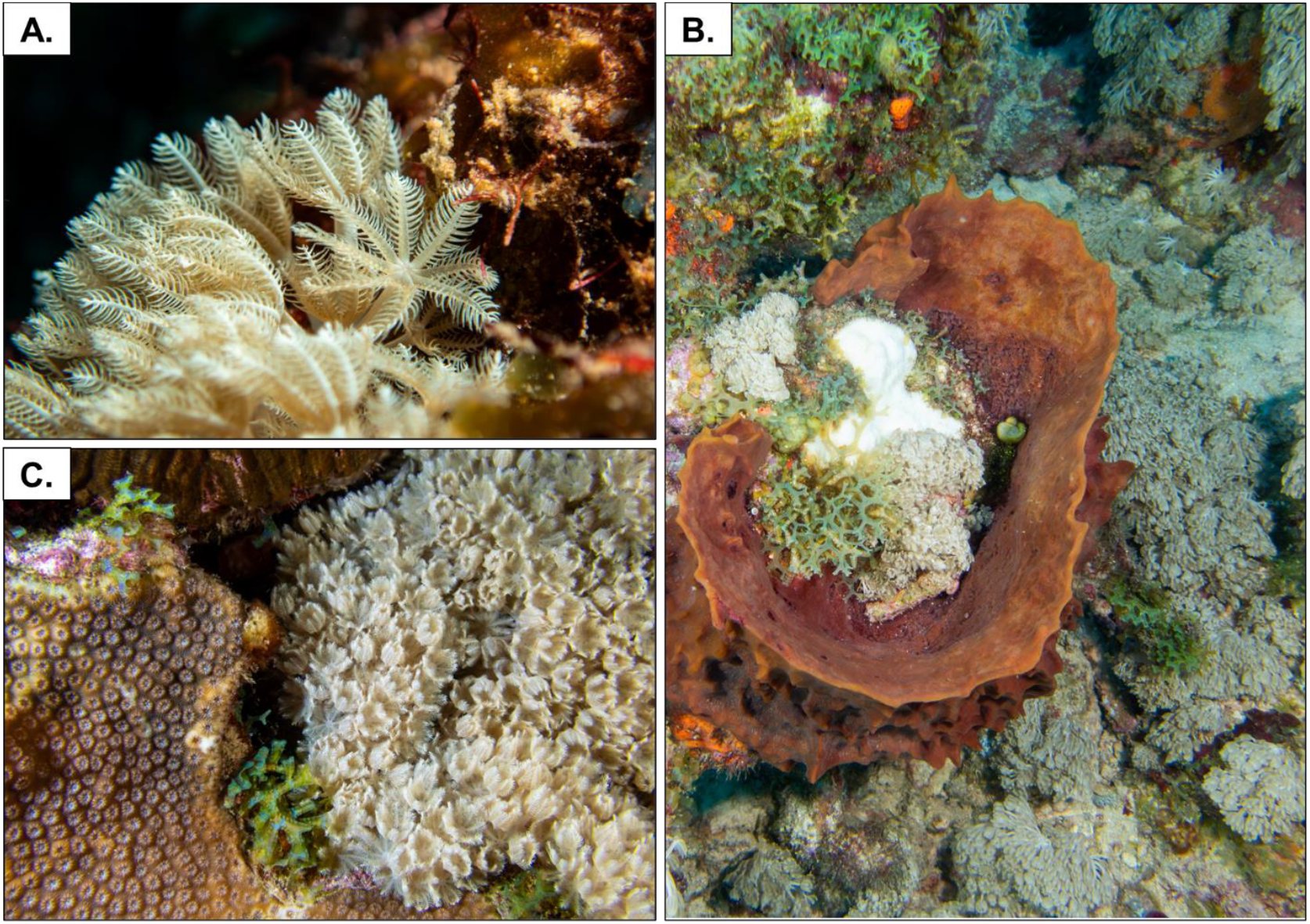
Field images of *Xenia umbellata* on reefs in La Parguera, Puerto Rico. (A) Close-up view highlighting polyp morphology. (B) *X. umbellata* colonizing substrate surrounding and within a giant barrel sponge (*Xestospongia muta*). (C) *X. umbellata* encroaching on the stony coral *Orbicella faveolata*. All photos by Daniel A. Toledo-Rodriguez.

Invasion genomics provides an effective evolutionary framework for investigating the invasion process, from potential introduction pathways to the establishment, spread, adaptation, and population dynamics of the invader (Lee, 2002; McGaughran et al., 2024; North et al., 2021). For example, by characterizing an introduced species’ genome, investigators can identify signatures of invasiveness, such as standing genetic variation (e.g., heterozygosity or admixture) or structural variation (e.g., gene duplications), which may facilitate rapid adaptation to the novel environment (Hahn & Rieseberg, 2017; Makino & Kawata, 2019; McGaughran et al., 2024; O’Donnell et al., 2014; Wu et al., 2019). Such genomic knowledge can help predict invasive potential, forecast invasion success, and support management priority setting in efforts to impede spread (McGaughran et al., 2024). Moreover, genomic resources generated during the early phases of an invasion provide critical baseline data for reconstructing invasion histories and identifying the genomic mechanisms of adaptation underlying the spread. While this information is typically generated only for the invading metazoan, it is equally important to characterize its associated microbial symbionts, which may influence the success and adaptability of the invader.

*Xenia umbellata*, like all cnidarians, harbors diverse microbial symbionts inclusive of dinoflagellates (Family Symbiodiniaceae), bacteria, archaea, fungi, and viruses, collectively referred to as the holobiont (Stévenne et al., 2021). The acquisition or loss of symbiotic partners can strongly influence host physiology by conferring new traits that may alter holobiont ecology and fitness (Bordenstein & Theis, 2015; Hussa & Goodrich-Blair, 2013; Pita et al., 2016). For example, in cnidarians, the identity of associated Symbiodiniaceae has been shown to influence host resilience to abiotic and biotic stressors, emphasizing the role of microbial symbionts in driving environmental adaptation and, ultimately, invasion success (Wang et al., 2023; Newkirk et al., 2020; Stévenne et al., 2021). Therefore, it is imperative to investigate the microbial symbionts of invaders and track changes in their communities as factors shaping invasion outcomes. In addition, these invasive microbial symbionts may directly affect native species by displacing resident symbionts or acting parasitically (Bojko et al., 2021). Considering both the host genome and its microbial symbionts as an integrated “hologenome” (Bordenstein & Theis, 2015) provides a powerful framework for understanding how invasions are mediated at the genomic and ecological levels and supports the need for hologenomic resources to guide future studies and management strategies.

Here, we describe the production of early-phase hologenomic resources for *Xenia umbellata* invading Puerto Rico’s coastal waters. Using deep metagenomic sequencing, we generated the first high-quality draft genome of *X. umbellata* and characterized the taxonomy of its associated Symbiodiniaceae. These resources were derived from a type specimen collected within six months of the initial report of the invasion. By integrating host and symbiont data, this study provides the first hologenomic reference for *X. umbellata*, establishing a foundation for both scientific investigation and management applications.

## 2. Data description

### 2.2 Sampling, DNA extraction, and Sequencing

Several polyps from a *Xenia umbellata* patch located in the La Parguera Natural Reserve at a depth of 21.6 m were carefully collected with tweezers on SCUBA and placed in a 50 mL tube in March 2024 (Table 1). The tissue sample in the 50 mL tube containing ambient seawater was then transferred to the lab on ice. Once in the lab, the ambient seawater was removed from the 50 mL tube. The polyps in the original 50 mL tube were then immediately placed in -80 ºC for storage until DNA extraction. DNA extraction was performed using the ZymoBIOMICS DNA/RNA Miniprep kit (Zymo Research) with minor modifications to the provider’s protocols (Veglia and Watkins 2025). Extracted DNA was sequenced on the Illumina NovaSeq 6000 platform using 150 bp paired-end reads, following library preparation with the Illumina DNA PCR-Free Prep kit. Sequencing was performed with a target output of 40 Gb, corresponding to approximately 133 million paired-end 150 bp reads (∼266 million total reads). Sample quality was assessed using Tapestation and Nanodrop, and library quality was evaluated using Tapestation and qPCR. DNA input met provider requirements: ≥10 ng/μL concentration, ≥0.1 μg total quantity, 260/280 ratio of 1.5-2.2, and DIN value between 6.0-10.0.

**Table 1.** MIxS data description for the *Xenia umbellata* collected from Puerto Rico coral reefs.

| Item | Definition |
| --- | --- |
| **General feature of classification** |  |
| Classification | Eukaryota; Opisthokonta; Metazoa; Eumetazoa; Cnidaria; Anthozoa; Octocorallia; Alcyonacea; Xeniidae; *Xenia*; *Xenia umbellata* |
| Investigation type | Eukaryote draft genome assembly |
| Project name | *Xenia umbellata* early Puerto Rico invasion hologenomic resources |
| Environment | Coral reef |
| Geographic location | Caribbean Sea; Puerto Rico; Lajas, La Parguera Natural Reserve; UR2 |
| Latitude, longitude | 17.895780 °N, -66.973600 °W |
| Collection date | 3/20/2024 |
| Environment properties | Insular shelf edge: spur and groove |
| Depth | 21.6 m |
| Collector | Daniel A. Toledo-Rodriguez |
| **Sequencing** |  |
| Sequencing method | Illumina NovaSeq 6000; paired-end reads (2x150) |
| Assembly method | *De novo* assembly |
| Program | SPAdes v4.0.0 |
| Finishing strategy | Scaffolding with RagTag v2.1.0 |
| **Accessibility** |  |
| DDBJ/ENA/GenBank | SAMN50581036; PRJNA1304975; GCA_021976095.1 |

### 2.2 Sequence processing and assembly

Sequencing resulted in 278,081,250 raw reads with 96% of bases having a phred score >20. Raw reads were then processed and cleaned with the program fastp (v0.23.2; Chen et al., 2018) resulting 272,355,896 high quality cleaned reads. High quality reads were assembled with the program SPAdes (v4.0.0; Prjibelski et al., 2020) using the metaSPAdes algorithm (Nurk et al., 2017) producing 1,092,378 scaffolds. The metagenome assembly provided a peak into the *X. umbellata* hologenome containing scaffolds sources from the xeniid metazoan as well as all its microbial symbionts inclusive of the endosymbiotic dinoflagellates within Family Symbiodiniaceae.

### 2.3 *Xenia umbellata* Genome Assembly, Assessment, and Annotation

To extract all xeniid scaffolds from the metagenome assembly, BLASTn (Camacho et al., 2009) was used to align contigs against a chromosome-level genome assembly of *Ovabunda* sp. (Hu et al., 2020). The genome assembly was originally labeled as *Xenia* sp., however it was later confirmed to be *Ovabunda* sp. (pers. comm. Catherine McFadden), a closely related genus. All sequences exhibiting >95% nucleotide identity and alignment lengths >100 bp were retained as putative xeniid scaffolds. On the remaining sequences, an additional scan for anthozoan-like scaffolds was performed using the program CAT (von Meijenfeldt et al., 2019) and the CAT_nr database (v20241212). All likely xeniid sequences (n=132,376), identified through BLASTn and/or CAT, were pooled into a single fasta file. The program RagTag (v2.1.0; Alonge et al., 2022) was then used to further scaffold these sequences using the chromosome-level *Ovabunda* genome assembly (Hu et al., 2020) as a reference. Next, the scaffolded assembly was then assessed for any contaminants (e.g., mitochondrial sequences, non-target organism sequences) to be removed using NCBI’s Foreign Contamination Screen (Astashyn et al., 2024). Finally, identified contaminants were removed and length filtering was then performed using the ‘clean’ function of the program funannotate (v1.8.17;Palmer & Stajich, 2020) resulting in a final assembly of 27,739 sequences with lengths greater than 500 nucleotides. Assembly quality was assessed using QUAST (v5.2.0; Mikheenko et al., 2023) revealing that the final assembly had a total length of 151,140,580 bp and an N50 of 6,477,837 bp (Figure 2). Genome completeness was assessed using BUSCO (v5.8.0; Manni et al., 2021) with the anthozoan lineage database (anthozoa_odb12.2025-07-01). The analysis revealed that 91.4% of BUSCOs were complete, comprising 89.9% single-copy and 1.5% duplicated genes. An additional 3.9% were fragmented and 4.7% were missing. Taken together, the QUAST and BUSCO results indicate high contiguity and completeness of the *Xenia umbellata* genome assembly. Next, we used the stats.sh program within the BBMAP tool kit (v39.15; Bushnell, 2014) to assess base content of the genome assembly. The genome assembly exhibited an overall base composition of 32.60% adenine (A), 32.82% thymine (T), 17.29% cytosine (C), and 17.29% guanine (G), with a GC content of 34.58% and 1.84% ambiguous bases (Ns), likely introduced during scaffolding and is expected for a draft genome assembled from short-read data. Further sequencing or long-read integration would likely improve contiguity, but this assembly represents a high-quality and biologically informative first reference genome for *Xenia umbellata*.

**Figure 2.**
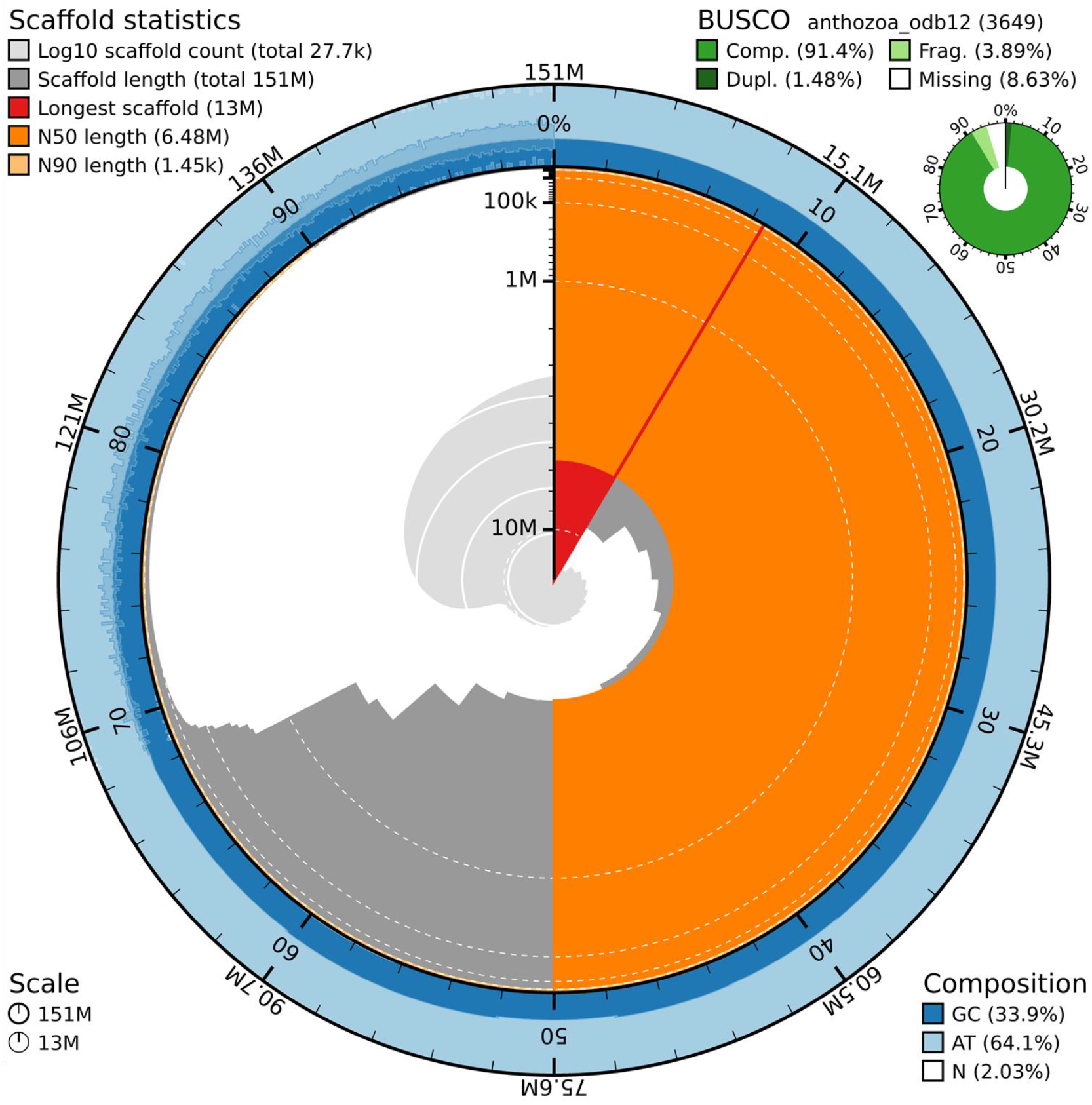
BlobToolKit (v4.4.5; Challis et al., 2020) snail plot illustrating scaffold metrics, completeness, and nucleotide composition of the draft genome assembly. The assembly comprises 27,739 scaffolds with a total length of 151.14 Mbp, a maximum scaffold size of 13 Mbp, an N50 of 6.48 Mbp, and an N90 of 1.45 kbp. BUSCO analysis against the anthozoa_odb12 dataset (n = 3,649 genes) recovered 91.4% complete (1.48% duplicated), 3.89% fragmented, and 8.63% missing orthologs, indicating high completeness. The genome has a GC content of 33.9%, AT content of 64.1%, and 2.03% ambiguous bases.

Prior to annotation, repetitive elements in the *X. umbellata* assembly were identified and classified *de novo* using RepeatModeler (v 2.0.7; Flynn et al., 2020). Identified repeats were then quantified and soft-masked assembly-wide with RepeatMasker (v4.2.1; Smit et al., 2013–2015) using both the custom library and the Dfam/RepBase databases as references. The *Xenia umbellata* genome was then annotated with the funannotate (v1.8.17) pipeline. Genes prediction was done with the *ab initio* predictors AUGUSTUS (v3.5.0; Stanke et al., 2006), GeneMark-ES (v4.71; Ter-Hovhannisyan et al., 2008), SNAP (v2006-07-28; Korf, 2004), and GlimmerHMM (v3.0.4; Majoros et al., 2004) and protein homology evidence generated by aligning NCBI RefSeq invertebrate proteins (downloaded July 2025) to the soft-masked genome with DIAMOND (Buchfink et al., 2015) and Exonerate (Slater & Birney, 2005). Consensus gene models were generated with EVidenceModeler (Haas et al., 2008), and functional annotation incorporated InterProScan (Jones et al., 2014), Pfam (Mistry et al., 2021), UniProt (v2025_03), MEROPS (v12.5; Rawlings et al., 2018), and dbCAN (v13; Zheng et al., 2023).

Annotation of the *Xenia umbellata* genome with Funannotate predicted 21,596 protein-coding genes across 20,844 mRNAs and 752 tRNAs, with an average gene length of ∼3.1 kb and an average protein length of 399 aa. The annotation comprised 142,785 exons (16,289 multi-exon and 4,555 single-exon transcripts). Functional annotation assigned putative functions to: 11,909 genes with GO terms, 14,275 InterProScan annotations, 11,452 Pfam domains, 692 MEROPS proteases, and 197 CAZymes. In addition, 1,143 genes were assigned common names through similarity to UniProt proteins.

### 2.4 Pairwise Genome Comparison with a Closely Related *Ovabunda* Species

Pairwise genome comparisons using PyANI-plus (ANIb method; v0.0.1; Pritchard et al., 2015) yielded an Average Nucleotide Identity (ANI) of 93.2% (*X. umbellata* aligned to *Ovabunda* sp.) and 92.1% (*Ovabunda* sp. aligned to *X. umbellata*), with a mean ANI of 92.7%. Alignment coverage was asymmetric, with 70.8% of the *X. umbellata* draft genome (107,045,860 bp) aligning to *Ovabunda* sp. and 63.6% of the *Ovabunda sp*. assembly (141,702,152 bp) aligning to *X. umbellata*. Transformed ANI (tANI) values were 0.415 and 0.535, respectively (mean = 0.475), consistent with substantial genomic divergence. These results indicate that while a large portion of the genome (∼64–71%) is shared at ∼93% identity, considerable lineage-specific sequence divergence remains, consistent with intergeneric genomic differentiation. Next, we calculated the estimated *X. umbellata* genome size using the program GenomeScope2 (v2.0; Ranallo-Benavidez et al., 2020). The kmer histogram file used for GenomeScope2 analyses was generated with the program jellyfish (v2.3.1; Marçais & Kingsford, 2011) with a “-m” equal to 21. GenomeScope analyses using 21-mer frequencies estimated the *X. umbellata* genome at ∼171.7 (171.6-171.9) Mbp (R^2^=93.15%). The estimated 171.7 Mbp genome size is 25.3 Mbp shorter than the calculated genome size for the related *Ovabunda* sp. genome. This observed difference is likely driven by repeat content (the “repeatome”), as GenomeScope estimated that the *X. umbellata* genome is approximately 38.1% repetitive (≈ 66 Mbp). In contrast, the *Ovabunda* sp. genome is 46.2% repetitive (≈ 91 Mbp) (Hu et al., 2020).

Furthermore, repeat region analyses identified 27.8% of the *X. umbellata* assembly (∼42.1 Mbp) as repetitive (26.97% interspersed repeats), dominated by unclassified elements (20.6%). Classified transposable elements comprised 6.4% of the genome (LINEs 2.6%, LTRs 1.5%, DNA transposons 2.3%). Transposable element expansions are a major driver of genomic divergence between species and have been reported to facilitate genome evolution in diverse eukaryotes (Castro et al., 2024; Pluess et al., 2016; Shah et al., 2020). Future efforts should build on this baseline characterization of the *X. umbellata* repeatome to assess changes in element abundance and diversity that may signal genomic adaptations to novel environments during its continued spread (Lee & Wang, 2018; Mérel et al., 2021).

### 2.5 *X. umbellata* Genome Heterozygosity: A Clue of Invasion Origin?

Genome-wide heterozygosity is hypothesized to provide insight into the adaptive and invasive potential of species (Kołodziejczyk et al., 2025). For example, the marbled crayfish (*Procambarus virginalis*), an emerging invasive species, possesses a triploid genome with high heterozygosity that is thought to facilitate its ecological success and spread (Gutekunst et al., 2018). In this context, establishing genome-wide heterozygosity for a specimen of *X. umbellata* collected during the early phase of an invasion provides a useful baseline for future comparisons, offers preliminary insight into adaptive capacity, and may help infer source populations: high diversity could indicate a wild origin, whereas reduced diversity might reflect bottlenecks associated with aquaculture or the aquarium trade. GenomeScope2 calculated the heterozygosity of the *X. umbellata* early-invasion genome to be approximately ∼1.3%. This value falls within the range that has been previously reported for cnidarians (0.79-1.96%; Locatelli & Baums, 2024; Shinzato et al., 2021; Stephens et al., 2022; Young et al., 2024; Yu et al., 2022) and is slightly higher than values reported for octocorals (0.73-1.2%; Ip et al., 2023; Ledoux et al., 2025). While this is a measurement of a single individual, it preliminarily suggests that the *X. umbellata* population introduced to Puerto Rican reefs may retain relatively high genomic diversity, consistent with high adaptive potential and a wild origin.

### 2.6 Characterization of *Xenia umbellata* dinoflagellate symbionts

An additional goal of this study was to provide genomic information/resources for the dinoflagellate symbionts (Family Symbiodiniaceae) associated with *Xenia umbellata* in Puerto Rico. Having this baseline knowledge is critical for tracking holobiont adaptation to the region via symbiont switching throughout *X. umbellata*’s continued spread (Creed et al., 2022; Sørensen et al., 2021). To conservatively identify Symbiodiniaceae scaffolds from the metagenome, we first aligned non-xeniid sequences to a Symbiodiniaceae reference database containing all publicly available genomes on NCBI (as of March 2025) using minimap2. Candidate scaffolds were then validated by re-alignment with BLASTn to the same database, with non-aligning sequences removed. Validated scaffolds were assigned genus-level taxonomy according to the best-matching reference genome. In total, 555,596 scaffolds, ranging in length from 200 to 25,187 bp, were identified as Symbiodiniaceae, with nucleotide similarities ranging from 95% to 100%. Of which, 555,520 (99.9% of scaffolds) aligned to the representative genomes from genus *Durusdinium*, suggesting the co-invading symbiont belongs to genus *Durusdinium*. Previous work reported *Xenia umbellata* in symbiosis with *Durusdinium*, but interestingly only at one shallow site in Ras Mohammed in the Red Sea, which was the only one of ten Red Sea sites where this association was observed (Osman et al., 2020). Our observation of *Durusdinium* as the dominant symbiont in this Puerto Rican individual suggests that *X. umbellata* has retained its original symbiosis during invasion. This finding provides critical baseline knowledge of the *X. umbellata*–*Durusdinium* association, enabling future efforts to track holobiont adaptation and monitor the spread of invasive symbionts.

## Acknowledgments

We thank Dr. Catherine S. McFadden for her continuous consultation on xeniids, Dr. Nilda Jimenez-Marrero (DNER-PR), Dr. Hector Ruiz-Torres (HJR Reefscaping) and the Department of Marine Sciences, UPRM, for providing field and logistical support. We acknowledge the Department of Natural and Environmental Resources for granting the sampling permit (O-VS-PVS15-SJ-01444-14032024) used in this study.

## Data accessibility

Raw reads can be accessed through the NBCI Sequence Read Archive (SRR34963192). This Genome Assembly project has been deposited at DDBJ/ENA/GenBank under the accession GCA_021976095.1. Data sources are linked to the NCBI BioSample and BioProject numbers SAMN50581036 and PRJNA1304975. All Symbiodiniaceae sequences and metadata are available for download on Zenodo (10.5281/zenodo.16849361)

## References

Alonge, M., Lebeigle, L., Kirsche, M., Jenike, K., Ou, S., Aganezov, S., Wang, X., Lippman, Z. B., Schatz, M. C., & Soyk, S. (2022). Automated assembly scaffolding using RagTag elevates a new tomato system for high-throughput genome editing. Genome Biology, 23(1), 258. 10.1186/s13059-022-02823-7

Astashyn, A., Tvedte, E. S., Sweeney, D., Sapojnikov, V., Bouk, N., Joukov, V., Mozes, E., Strope, P. K., Sylla, P. M., Wagner, L., Bidwell, S. L., Brown, L. C., Clark, K., Davis, E. W., Smith-White, B., Hlavina, W., Pruitt, K. D., Schneider, V. A., & Murphy, T. D. (2024). Rapid and sensitive detection of genome contamination at scale with FCS-GX. Genome Biology, 25(1), 60. 10.1186/s13059-024-03198-7

Bojko, J., Burgess, A. L., Baker, A. G., & Orr, C. H. (2021). Invasive Non-Native Crustacean Symbionts: Diversity and Impact. Journal of Invertebrate Pathology, 186, 107482. 10.1016/j.jip.2020.107482

Bordenstein, S. R., & Theis, K. R. (2015). Host Biology in Light of the Microbiome: Ten Principles of Holobionts and Hologenomes. PLoS Biology, 13(8), e1002226. 10.1371/journal.pbio.1002226

Buchfink, B., Xie, C., & Huson, D. H. (2015). Fast and sensitive protein alignment using DIAMOND. Nature Methods, 12(1), Article 1. 10.1038/nmeth.3176

Bushnell, B. (2014, March 17). BBMap: A Fast, Accurate, Splice-Aware Aligner. https://www.semanticscholar.org/paper/BBMap%3A-A-Fast%2C-Accurate%2C-Splice-Aware-Aligner-Bushnell/f64dd54444a724574deb7710888091350eebb2b9

Camacho, C., Coulouris, G., Avagyan, V., Ma, N., Papadopoulos, J., Bealer, K., & Madden, T. L. (2009). BLAST+: Architecture and applications. BMC Bioinformatics, 10, 421. 10.1186/1471-2105-10-421

Castro, N., Vilela, B., Mata-Sucre, Y., Marques, A., Gagnon, E., Lewis, G. P., Costa, L., & Souza, G. (n.d.). Repeatome evolution across space and time: Unravelling repeats dynamics in the plant genus Erythrostemon Klotzsch (Leguminosae Juss). Molecular Ecology, n/a(n/a), e17510. 10.1111/mec.17510

Challis, R., Richards, E., Rajan, J., Cochrane, G., & Blaxter, M. (2020). BlobToolKit: Interactive quality assessment of genome assemblies. G3: Genes, Genomes, Genetics, 10(4), 1361–1374. 10.1534/g3.119.400908

Chen, S., Zhou, Y., Chen, Y., & Gu, J. (2018). fastp: An ultra-fast all-in-one FASTQ preprocessor. Bioinformatics, 34(17), i884–i890. 10.1093/bioinformatics/bty560

Creed, R. P., Brown, B. L., & Skelton, J. (2022). The potential impacts of invasions on native symbionts. Ecology, 103(8), e3726. 10.1002/ecy.3726

Gutekunst, J., Andriantsoa, R., Falckenhayn, C., Hanna, K., Stein, W., Rasamy, J., & Lyko, F. (2018). Clonal genome evolution and rapid invasive spread of the marbled crayfish. Nature Ecology & Evolution, 2(3), 567–573. 10.1038/s41559-018-0467-9

Haas, B. J., Salzberg, S. L., Zhu, W., Pertea, M., Allen, J. E., Orvis, J., White, O., Buell, C. R., & Wortman, J. R. (2008). Automated eukaryotic gene structure annotation using EVidenceModeler and the Program to Assemble Spliced Alignments. Genome Biology, 9(1), R7. 10.1186/gb-2008-9-1-r7

Hahn, M. A., & Rieseberg, L. H. (2017). Genetic admixture and heterosis may enhance the invasiveness of common ragweed. Evolutionary Applications, 10(3), 241–250. 10.1111/eva.12445

Halász, A., Mcfadden, C. S., Toonen, R., & Benayahu, Y. (2019). Re-description of type material of Xenia Lamarck, 1816 (Octocorallia: Xeniidae). Zootaxa, 4652(2), 201–239. 10.11646/zootaxa.4652.2.1

Hu, M., Zheng, X., Fan, C.-M., & Zheng, Y. (2020). Lineage dynamics of the endosymbiotic cell type in the soft coral Xenia. Nature, 582(7813), 534–538. 10.1038/s41586-020-2385-7

Hussa, E. A., & Goodrich-Blair, H. (2013). It Takes a Village: Ecological and Fitness Impacts of Multipartite Mutualism. Annual Review of Microbiology, 67(1), 161–178. 10.1146/annurev-micro-092412-155723

Ip, J. C.-H., Ho, M.-H., Chan, B. K. K., & Qiu, J.-W. (2023). A draft genome assembly of reef-building octocoral Heliopora coerulea. Scientific Data, 10(1), 381. 10.1038/s41597-023-02291-z

Jones, P., Binns, D., Chang, H.-Y., Fraser, M., Li, W., McAnulla, C., McWilliam, H., Maslen, J., Mitchell, A., Nuka, G., Pesseat, S., Quinn, A. F., Sangrador-Vegas, A., Scheremetjew, M., Yong, S.-Y., Lopez, R., & Hunter, S. (2014). InterProScan 5: Genome-scale protein function classification. Bioinformatics, 30(9), 1236–1240. 10.1093/bioinformatics/btu031

Kołodziejczyk, J., Fijarczyk, A., Porth, I., Robakowski, P., Vella, N., Vella, A., Kloch, A., & Biedrzycka, A. (2025). Genomic investigations of successful invasions: The picture emerging from recent studies. Biological Reviews of the Cambridge Philosophical Society, 100(3), 1396–1418. 10.1111/brv.70005

Korf, I. (2004). Gene finding in novel genomes. BMC Bioinformatics, 5(1), 59. 10.1186/1471-2105-5-59

Ledoux, J.B., Gomez-Garrido, J., Cruz, F., Camara Ferreira, F., Matos, A., Sarropoulou, X., Ramirez-Calero, S., Aurelle, D., Lopez-Sendino, P., Grayson, N. E., Moore, B. S., Antunes, A., Aguilera, L., Gut, M., Salces-Ortiz, J., Fernández, R., Linares, C., Garrabou, J., & Alioto, T. (2025). Chromosome-Level Genome Assembly and Annotation of Corallium rubrum: A Mediterranean Coral Threatened by Overharvesting and Climate Change. Genome Biology and Evolution, 17(2), evae253. 10.1093/gbe/evae253

Lee, C. E. (2002). Evolutionary genetics of invasive species. Trends in Ecology & Evolution, 17(8), 386–391. 10.1016/S0169-5347(02)02554-5

Lee, C.C., & Wang, J. (2018). Rapid Expansion of a Highly Germline-Expressed Mariner Element Acquired by Horizontal Transfer in the Fire Ant Genome. Genome Biology and Evolution, 10(12), 3262–3278. 10.1093/gbe/evy220

Lineage-specific symbionts mediate differential coral responses to thermal stress | Microbiome | Full Text. (n.d.). Retrieved September 4, 2025, from https://microbiomejournal.biomedcentral.com/articles/10.1186/s40168-023-01653-4

Locatelli, N. S., & Baums, I. B. (2024). Genomes of the Caribbean reef-building corals Colpophyllia natans, Dendrogyra cylindrus, and Siderastrea siderea. bioRxiv, 2024.08.21.608299. 10.1101/2024.08.21.608299

Majoros, W. H., Pertea, M., & Salzberg, S. L. (2004). TigrScan and GlimmerHMM: Two open source ab initio eukaryotic gene-finders. Bioinformatics, 20(16), 2878–2879. 10.1093/bioinformatics/bth315

Makino, T., & Kawata, M. (2019). Invasive invertebrates associated with highly duplicated gene content. Molecular Ecology, 28(7), 1652–1663. 10.1111/mec.15019

Manni, M., Berkeley, M. R., Seppey, M., Simao, F. A., & Zdobnov, E. M. (2021). BUSCO update: Novel and streamlined workflows along with broader and deeper phylogenetic coverage for scoring of eukaryotic, prokaryotic, and viral genomes (No. arXiv:2106.11799). arXiv. 10.48550/arXiv.2106.11799

Marçais, G., & Kingsford, C. (2011). A fast, lock-free approach for efficient parallel counting of occurrences of k-mers. Bioinformatics, 27(6), 764–770. 10.1093/bioinformatics/btr011

McGaughran, A., Dhami, M. K., Parvizi, E., Vaughan, A. L., Gleeson, D. M., Hodgins, K. A., Rollins, L. A., Tepolt, C. K., Turner, K. G., Atsawawaranunt, K., Battlay, P., Congrains, C., Crottini, A., Dennis, T. P. W., Lange, C., Liu, X. P., Matheson, P., North, H. L., Popovic, I., … Wilson, J. (2024). Genomic Tools in Biological Invasions: Current State and Future Frontiers. Genome Biology and Evolution, 16(1), evad230. 10.1093/gbe/evad230

Mérel, V., Gibert, P., Buch, I., Rodriguez Rada, V., Estoup, A., Gautier, M., Fablet, M., Boulesteix, M., & Vieira, C. (2021). The Worldwide Invasion of Drosophila suzukii Is Accompanied by a Large Increase of Transposable Element Load and a Small Number of Putatively Adaptive Insertions. Molecular Biology and Evolution, 38(10), 4252–4267. 10.1093/molbev/msab155

Mezger, S. D., Klinke, A., Tilstra, A., El-Khaled, Y. C., Thobor, B., & Wild, C. (2022). The widely distributed soft coral Xenia umbellata exhibits high resistance against phosphate enrichment and temperature increase. Scientific Reports, 12, 22135. 10.1038/s41598-022-26082-9

Mikheenko, A., Saveliev, V., Hirsch, P., & Gurevich, A. (2023). WebQUAST: Online evaluation of genome assemblies. Nucleic Acids Research, 51(W1), W601–W606. 10.1093/nar/gkad406

Mistry, J., Chuguransky, S., Williams, L., Qureshi, M., Salazar, G. A., Sonnhammer, E. L. L., Tosatto, S. C. E., Paladin, L., Raj, S., Richardson, L. J., Finn, R. D., & Bateman, A. (2021). Pfam: The protein families database in 2021. Nucleic Acids Research, 49(D1), D412–D419. 10.1093/nar/gkaa913

Nadir, E., Lotan, T., & Benayahu, Y. (2023). Xenia umbellata (Octocorallia): A novel model organism for studying octocoral regeneration ability. Frontiers in Marine Science, 10. 10.3389/fmars.2023.1021679

Newkirk, C. R., Frazer, T. K., Martindale, M. Q., & Schnitzler, C. E. (2020). Adaptation to Bleaching: Are Thermotolerant Symbiodiniaceae Strains More Successful Than Other Strains Under Elevated Temperatures in a Model Symbiotic Cnidarian? Frontiers in Microbiology, 11. 10.3389/fmicb.2020.00822

North, H. L., McGaughran, A., & Jiggins, C. D. (2021). Insights into invasive species from whole-genome resequencing. Molecular Ecology, 30(23), 6289–6308. 10.1111/mec.15999

Nurk, S., Meleshko, D., Korobeynikov, A., & Pevzner, P. A. (2017). metaSPAdes: A new versatile metagenomic assembler. Genome Research, 27(5), 824–834. 10.1101/gr.213959.116

O’Donnell, D. R., Parigi, A., Fish, J. A., Dworkin, I., & Wagner, A. P. (2014). The Roles of Standing Genetic Variation and Evolutionary History in Determining the Evolvability of Anti-Predator Strategies. PLoS ONE, 9(6), e100163. 10.1371/journal.pone.0100163

Osman, E. O., Suggett, D. J., Voolstra, C. R., Pettay, D. T., Clark, D. R., Pogoreutz, C., Sampayo, E. M., Warner, M. E., & Smith, D. J. (2020). Coral microbiome composition along the northern Red Sea suggests high plasticity of bacterial and specificity of endosymbiotic dinoflagellate communities. Microbiome, 8(1), Article 1. 10.1186/s40168-019-0776-5

Palmer, J. M., & Stajich, J. (2020). Funannotate v1.8.1: Eukaryotic genome annotation [Computer software]. Zenodo. 10.5281/zenodo.4054262

Pita, L., Fraune, S., & Hentschel, U. (2016). Emerging Sponge Models of Animal-Microbe Symbioses. Frontiers in Microbiology, 7. https://www.frontiersin.org/articles/10.3389/fmicb.2016.02102

Pluess, A. R., Frank, A., Heiri, C., Lalagüe, H., Vendramin, G. G., & Oddou-Muratorio, S. (2016). Genome–environment association study suggests local adaptation to climate at the regional scale in Fagus sylvatica. New Phytologist, 210(2), 589–601. 10.1111/nph.13809

Pritchard, L., Glover, R. H., Humphris, S., Elphinstone, J. G., & Toth, I. K. (2015). Genomics and taxonomy in diagnostics for food security: Soft-rotting enterobacterial plant pathogens. Analytical Methods, 8(1), 12–24. 10.1039/C5AY02550H

Prjibelski, A., Antipov, D., Meleshko, D., Lapidus, A., & Korobeynikov, A. (2020). Using SPAdes De Novo Assembler. Current Protocols in Bioinformatics, 70(1), e102. 10.1002/cpbi.102

Ranallo-Benavidez, T. R., Jaron, K. S., & Schatz, M. C. (2020). GenomeScope 2.0 and Smudgeplot for reference-free profiling of polyploid genomes. Nature Communications, 11(1), 1432. 10.1038/s41467-020-14998-3

Rawlings, N. D., Barrett, A. J., Thomas, P. D., Huang, X., Bateman, A., & Finn, R. D. (2018). The MEROPS database of proteolytic enzymes, their substrates and inhibitors in 2017 and a comparison with peptidases in the PANTHER database. Nucleic Acids Research, 46(D1), D624–D632. 10.1093/nar/gkx1134

Ruiz-Allais, J. P., Amaro, M. E., McFadden, C. S., Halász, A., & Benayahu, Y. (2014). The first incidence of an alien soft coral of the family Xeniidae in the Caribbean, an invasion in eastern Venezuelan coral communities. Coral Reefs, 33(2), 287–287. doi:10.1007/s00338-013-1122-1

Ruiz-Allais, J. P., Benayahu, Y., & Lasso-Alcalá, O. M. (2021). The invasive octocoral Unomia stolonifera (Alcyonacea, Xeniidae) is dominating the benthos in the Southeastern Caribbean Sea. doi:10.5281/ZENODO.4784709

Shah, A., Hoffman, J. I., & Schielzeth, H. (2020). Comparative Analysis of Genomic Repeat Content in Gomphocerine Grasshoppers Reveals Expansion of Satellite DNA and Helitrons in Species with Unusually Large Genomes. Genome Biology and Evolution, 12(7), 1180–1193. 10.1093/gbe/evaa119

Shinzato, C., Takeuchi, T., Yoshioka, Y., Tada, I., Kanda, M., Broussard, C., Iguchi, A., Kusakabe, M., Marin, F., Satoh, N., & Inoue, M. (2021). Whole-Genome Sequencing Highlights Conservative Genomic Strategies of a Stress-Tolerant, Long-Lived Scleractinian Coral, Porites australiensis Vaughan, 1918. Genome Biology and Evolution, 13(12), evab270. 10.1093/gbe/evab270

Slater, G. S. C., & Birney, E. (2005). Automated generation of heuristics for biological sequence comparison. BMC Bioinformatics, 6(1), 31. 10.1186/1471-2105-6-31

Sørensen, M. E. S., Wood, A. J., Cameron, D. D., & Brockhurst, M. A. (2021). Rapid compensatory evolution can rescue low fitness symbioses following partner switching. Current Biology, 31(17), 3721-3728.e4. 10.1016/j.cub.2021.06.034

Stanke, M., Keller, O., Gunduz, I., Hayes, A., Waack, S., & Morgenstern, B. (2006). AUGUSTUS: Ab initio prediction of alternative transcripts. Nucleic Acids Research, 34(suppl_2), W435–W439. 10.1093/nar/gkl200

Stephens, T. G., Lee, J., Jeong, Y., Yoon, H. S., Putnam, H. M., Majerová, E., & Bhattacharya, D. (2022). High-quality genome assembles from key Hawaiian coral species. GigaScience, 11, giac098. 10.1093/gigascience/giac098

Stévenne, C., Micha, M., Plumier, J.-C., & Roberty, S. (2021). Corals and Sponges Under the Light of the Holobiont Concept: How Microbiomes Underpin Our Understanding of Marine Ecosystems. Frontiers in Marine Science, 8. 10.3389/fmars.2021.698853

Ter-Hovhannisyan, V., Lomsadze, A., Chernoff, Y. O., & Borodovsky, M. (2008). Gene prediction in novel fungal genomes using an ab initio algorithm with unsupervised training. Genome Research, 18(12), 1979–1990. 10.1101/gr.081612.108

Toledo-Rodriguez, D. A., Veglia, A. J., Marrero, N. M. J., Gomez-Samot, J. M., McFadden, C. S., Weil, E., & Schizas, N. V. (2025). Shadows over Caribbean reefs: Occurrence of a new invasive soft coral species, Xenia umbellata, in southwest Puerto Rico. Coral Reefs, 44(4), 1439–1444. 10.1007/s00338-025-02670-5

Veglia, A. J., & Watkins, J. A. (2025). ZymoBIOMICS DNA/RNA Miniprep Kit – ViDaB Lab. protocols.io. 10.17504/protocols.io.6qpvrw41zlmk/v1

von Meijenfeldt, F. A. B., Arkhipova, K., Cambuy, D. D., Coutinho, F. H., & Dutilh, B. E. (2019). Robust taxonomic classification of uncharted microbial sequences and bins with CAT and BAT. Genome Biology, 20(1), 217. 10.1186/s13059-019-1817-x

Wang, C., Zheng, X., Kvitt, H., Sheng, H., Sun, D., Niu, G., Tchernov, D., & Shi, T. (2023). Lineage-specific symbionts mediate differential coral responses to thermal stress. Microbiome, 11, 211. 10.1186/s40168-023-01694-1

Wu, N., Zhang, S., Li, X., Cao, Y., Liu, X., Wang, Q., Liu, Q., Liu, H., Hu, X., Zhou, X. J., James, A. A., Zhang, Z., Huang, Y., & Zhan, S. (2019). Fall webworm genomes yield insights into rapid adaptation of invasive species. Nature Ecology & Evolution, 3(1), 105–115. 10.1038/s41559-018-0746-5

Young, B. D., Williamson, O. M., Kron, N. S., Andrade Rodriguez, N., Isma, L. M., MacKnight, N. J., Muller, E. M., Rosales, S. M., Sirotzke, S. M., Traylor-Knowles, N., Williams, S. D., & Studivan, M. S. (2024). Annotated genome and transcriptome of the endangered Caribbean mountainous star coral (Orbicella faveolata) using PacBio long-read sequencing. BMC Genomics, 25, 226. 10.1186/s12864-024-10092-w

Yu, Y., Nong, W., So, W. L., Xie, Y., Yip, H. Y., Haimovitz, J., Swale, T., Baker, D. M., Bendena, W. G., Chan, T. F., Chui, A. P. Y., Lau, K. F., Qian, P.-Y., Qiu, J.-W., Thibodeau, B., Xu, F., & Hui, J. H. L. (2022). Genome of elegance coral Catalaphyllia jardinei (Euphylliidae). Frontiers in Marine Science, 9. 10.3389/fmars.2022.991391

Zheng, J., Ge, Q., Yan, Y., Zhang, X., Huang, L., & Yin, Y. (2023). dbCAN3: Automated carbohydrate-active enzyme and substrate annotation. Nucleic Acids Research, 51(W1), W115–W121. 10.1093/nar/gkad328

